# SynapsEM: computer-assisted synapse morphometry

**DOI:** 10.1101/2020.08.28.271882

**Authors:** Shigeki Watanabe, M Wayne Davis, Grant F Kusick, Janet Iwasa, Erik M Jorgensen

## Abstract

The structural features of a synapse, in part, determine its output. Synapses are extremely small and tightly packed with vesicles and other organelles. Visualizing synaptic structure requires imaging by electron microscopy, and the features in micrographs must be quantified using morphometry. Three parameters are typically assessed from each specimen: 1) the sizes of individual vesicles and organelles, 2) the absolute number and densities of organelles, and 3) distances between organelles and key features at synapses such as active zone membranes and dense projections. For data to be valid, the analysis must be repeated from hundreds or thousands of images from several biological replicates, a daunting task. Here we report a custom computer program to analyze these features: SynapsEM. In short, we developed macros for ImageJ/Fiji to record x,y-coordinates of segmented structures; the coordinates are then exported as text files. Independent investigators can reload the images and text files to confirm or re-evaluate the segmentation using ImageJ. The Matlab program calculates and reports key synaptic parameters from the coordinates. Since the values are calculated, rather than measured from each micrograph, other parameters can be extracted in Matlab by additional scripting. Thus, this program can accelerate morphometry of synapses and promote a more comprehensive analysis of synaptic ultrastructure.

## 1 Introduction

Understanding the mechanisms of synaptic transmission requires detailed characterizations of synapses at the ultrastructural level. To release neurotransmitter, synaptic vesicles fuse at a specialized membrane domain of the presynaptic terminal called the active zone (Couteaux and Pécot-Dechavassine, 1970; Heuser et al., 1979). A subset of vesicles are docked, that is, in contact with the active zone membrane by morphology (Hammarlund et al., 2007; Imig et al., 2014; Schikorski and Stevens, 1997), and fuse in response to calcium influx (Heuser and Reese, 1981; Heuser et al., 1979). Following fusion, these vesicles are recycled locally via endocytosis and components sorted in an endosome to sustain synaptic transmission (Ceccarelli et al., 1972; Dittman and Ryan, 2009; Heuser and Reese, 1973; Kononenko and Haucke, 2015; Saheki and De Camilli, 2012; Watanabe et al., 2013a, 2013b, 2014). Unfortunately, the structures involved in synaptic membrane trafficking are extremely small. For example, synaptic vesicles are 30-50 nm in diameter, and a few hundred vesicles are clustered within a synaptic bouton (Schikorski and Stevens, 1997; Shepherd and Harris, 1998), which is only ∼0.5-1 µm in diameter. Moreover, a vesicle may move only a few nanometers to fully engage the active zone membrane during docking (Hammarlund et al., 2007; Imig et al., 2014), and this state is quite dynamic (Chang et al., 2018; Kusick et al., 2018). Given these dimensions, synaptic morphometry requires the resolution of electron microscopy.

Morphometry is typically performed on an electron micrograph of single synaptic profile from a 30-70 nm-thick section. Analyses from ∼ 200 synaptic profiles are then summed, and results compared between controls and experimental samples, such as mutants or drug-treated neurons. In each image, the following features are analyzed: the size of membrane-bound structures such as vesicles and other organelles, the number of these structures, and the distance of these structures from the active zone membrane, the plasma membrane, and if apparent, electron-dense cytomatrix (dense projection) that harbors calcium channels. The organelles are identified in serial electron micrographs, that is, they are ‘segmented’, and characterized manually by measuring the sizes of the organelles and distances of these structures to the relevant membranes. Thus, the process is labor-intensive, and typically does not provide a reliable record of annotated features.

To overcome these issues, we developed an analysis workflow, SynapsEM, that integrates ImageJ macros and Matlab scripts for the morphometry of synapses from electron micrographs (Figure 1). Specifically, the Matlab scripts first shuffle images from different conditions, which are pooled into a single folder (Figure 1A). This shuffling procedure reduces the potential bias during annotation. These images are imported into ImageJ as a sequence (Figure 1A). With the freehand line tool, the contours of the plasma membrane and the active zone membrane are traced, and their x,y-coordinates recorded in the ROI manager (Figure 1B). Then, the diameters of vesicles are traced using a straight line tool (Figure 1B inset), and the x,y-coordinates at the centroid of vesicles recorded. Additionally, endosomes can be traced with a freehand selection tool, and their positions recorded. Once all structures are annotated from an image, the values in the ROI are exported as a text file (Figure 1B, C). The text file can be imported back to ImageJ for re-evaluation by independent experimenters. This re-evaluation step is not required but independent confirmation of segmentation calls makes the annotation more accurate. When all images are analyzed, the resulting text files are unblinded and imported into Matlab (Figure 1C). The custom scripts then calculate the distances of membrane-bound structures to the active zone membrane as well as the plasma membrane. The numbers and diameters of these structures are also determined. These data are then compared between different conditions by the computer; the researcher remains blinded to individual images. The outputs of the scripts can be saved or exported to other programs for statistical analysis. Overall, this workflow expedites the analysis of synaptic ultrastructure, reduces the experimental biases associated with manual image annotation, and unifies the method of analysis across many labs.

**Figure 1.**
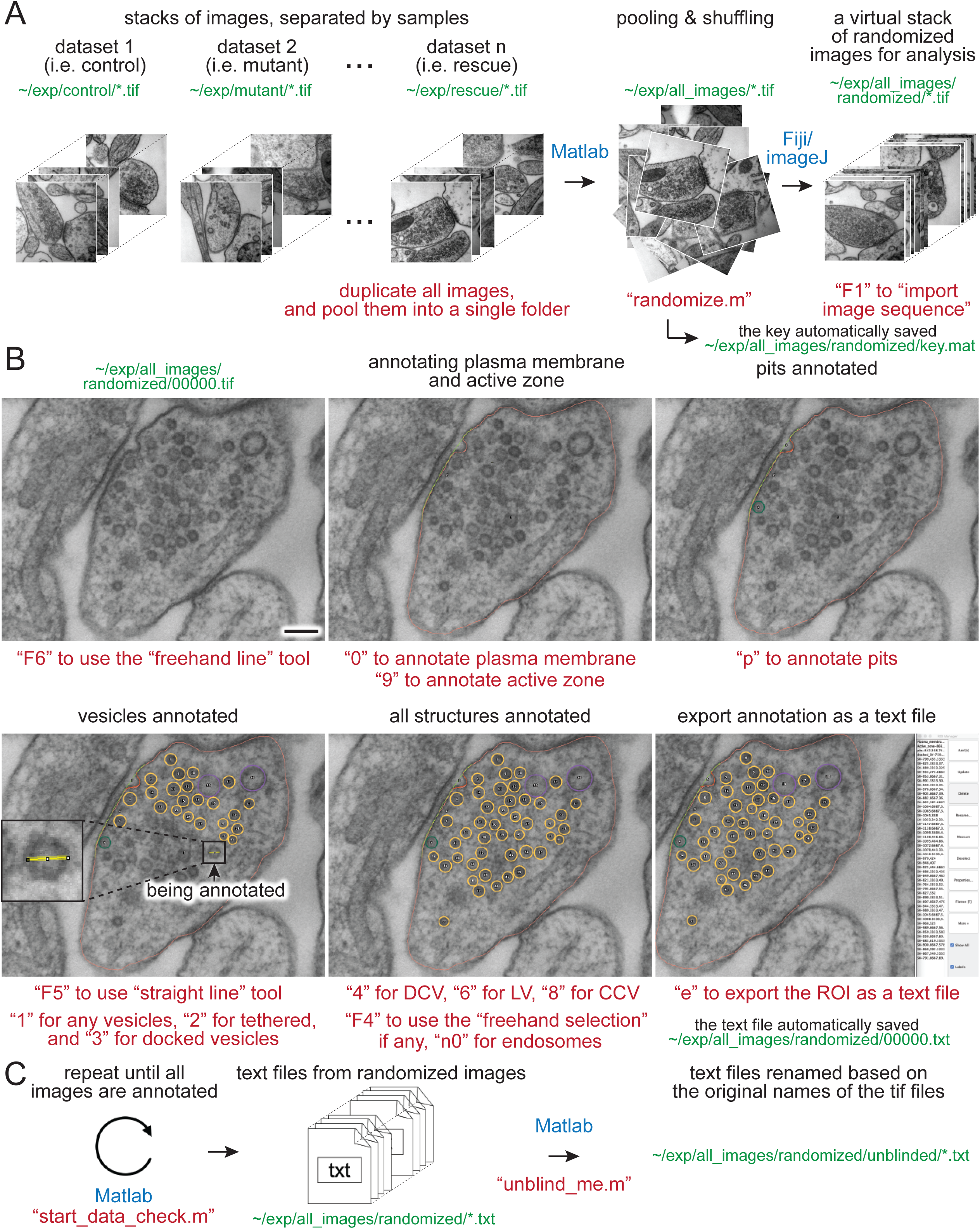
Synapse morphometry using synapsEM. (A) A schematic showing the image randomization procedure. In each experiment, images are collected from multiple samples (i.e. control, mutants, drug-treated). Ideally, image acquisition should also be done blinded. These images are pooled into the same folder and then randomized using the “randomize.m” Matlab script. Running this program creates a new folder, named “randomized”, and transfers the images into this folder with the randomized number assigned to each image. These images should be opened in ImageJ or Fiji as a virtual stack using the F1 key after installing the “synapsEM_analysis_macro.txt” macro. (B) A schematic showing the procedure for image annotation. After opening the images in ImageJ/Fiji, membranes can be traced using specific tools and hot keys (as denoted in red). Note that all structures annotated are recorded into the ROI manager. For vesicles, draw a line across the vesicle membrane as shown in the inset. After pressing a hot key corresponding to the structure (i.e. “1” for synaptic vesicle in this case), the macro draws a circle using the drawn line as a diameter for the vesicle. When the annotation is complete, the structures listed in the ROI manager can be exported by simply pressing “e” on the computer keyboard. (C) A schematic showing the final steps of the morphometry. After all images in the dataset are annotated, the resulting text files must be first checked for errors using the “start_data_check” script in Matlab. After corrections, the text files can be decoded based on the original names of the corresponding images using the “unblind_me” script. This script duplicates the text files, decode the copied files, and move them to the “unblinded” folder. The decoded text files can be further processed for analysis. The sentences in red, green, and blue indicate the commands, the directory of files, and the software used, respectively. Scale bars = 100 nm.

## Materials and equipment

The following materials are required for the procedures described in this manuscript.

- Electron micrographs, preferably acquired with the same camera setting on the same microscope for a set of experiments.
- A computer (no specific requirements as long as it meets the system requirement for the Matlab program).
- A computer keyboard with the numeric key pad.
- Matlab, MathWorks. (The scripts were originally written in Matlab 2008, and have been added to in versions up to Matlab 2020.)
- Matlab custom scripts (available from **https://github.com/shigekiwatanabe/SynapsEM**).
- Fiji (or ImageJ).
- Macros (available from **https://github.com/shigekiwatanabe/SynapsEM**).
- Maya for 3-D rendering (optional).
- Wacom tablet (cintiq 22HD), or other pen tablet (optional, but makes annotating many images easier).

## Methods

### 3.1 Randomizing images

Images from multiple samples should be analyzed in a batch to minimize the potential bias during the analysis. Shuffling removes any possibility of conscious or unconscious bias between samples, eliminating any variables in analysis apart from different samples themselves. Even with blinded, but not scrambled images, it is possible for the analysis to be skewed when they are analyzed in different batches: this can range from an obvious phenotype making it clear which condition a sample is, to simple day-to-day differences in segmentation, to a novice improving the accuracy of their segmentation as they work through more images in an experiment. For this purpose, all images for single experiments are pooled into one folder (Figure 1A). These images should be duplicates of the original images so as to keep the original data untouched. To ensure the original images are safe, the program asks whether the images should be duplicated when the “randomize” code is run in Matlab. Answering “yes” to this question will make copies of the images and randomize the duplicated images, leaving the original data untouched. In the popup window, select the directory (folder) that contains all images for an experiment. After the selection, another window pops up, prompting the user to select all images. Select all images to be randomized. At the end of the program, randomized images are found in the “randomized” folder, nested in the directory where the original images are (Figure 1A). The key is named “key.mat” and also saved in the “randomized” folder (Figure 1A). It is important that this file is kept safe until the analysis is complete.

### 3.2 Opening images

Morphometry is performed in Fiji. Start Fiji, and install the “synapsEM_analysis_macro.txt” by navigating to the plugin dropdown menu, clicking on “Macros” and then “Install”, and selecting the “synapsEM_analysis_macro.txt” file. For easy access to the file, it is highly recommended to store this macro in the ∼/Fiji/macros directory so it is readily accessible. To skip this installation procedure, the contents of “synapsEM_analysis_macro.txt” can be copied into the StartupMacros.fiji.ijm, which is found in the ∼/Fiji/macros directory, since all the macros in this file are activated as Fiji starts up.

To start the analysis, the randomized images should be opened as a stack (Figure 1A). Press the “F1” key to import the image sequence as a virtual stack – the pixel size information on tif files will be converted such that each pixel is one arbitrary unit when the “F1” hot key is used. If images are opened through “File, Import, Image sequence” or simply dragging a folder to Fiji, it is important to “set scale” through the “Analyze” drop-down menu and type in 1 for the “distance in pixels” and 1 for the “known distance”. This conversion of the pixel size can also be performed by pressing “F2”. When working with a large dataset, it is highly recommended that images are opened as a virtual stack.

### 3.3 Segmenting images

The eventual goal of segmentation is to determine size of vesicles and other membrane-bound organelles, numbers or density of these structures, and the distribution of these structures relative to the plasma membrane or active zone membrane. The macros are set up to record x,y-coordinates of the structures of interest and their size information in the ROI manager window. These macros are accessed through hot keys, as listed in Table 1 and 2. Three sets of line tools are used to trace different features at synapses. A straight line tool (the Fiji tool #1) is accessed with “F5” and used to annotate closed and near-uniformly circular structures like synaptic vesicles, large vesicles, and dense-core vesicles. A freehand selection tool (the Fiji tool #3, “F4”) is used to segment closed and irregularly-shaped structures such as endosomes. A freehand line tool (the Fiji tool #7, “F6”) is used to segment open-ended structures, such as plasma membranes, active zone membranes, and pits. When an appropriate tool in Fiji is selected (Table 1), objects in the micrograph can be segmented using hot keys listed in Table 2 (Figure 1). For open-ended membranes, trace the contour of the membranes as closely as possible using the freehand line tool. From each micrograph, one plasma membrane (“0”) and at least one active zone membrane (“9”) must be segmented. Since some synapses have multiple active zones (Figure 2A, B), judged based on the presence of postsynaptic density in the juxtaposed membranes, more than one active zone can be defined per micrograph. However, a single plasma membrane should be drawn in each image. In some synapses, presynaptic dense projections are prominent in the active zone (Watanabe et al., 2013a; Zhai and Bellen, 2004), and they can be traced using the same tool and added to the ROI manager by pressing “d” for dense projection and “r” for synaptic ribbon. If membranes are deflected inward, towards the cytoplasm, (Figure 1A-C, 2E,F), they can be traced as pits (“U”), although the exact nature of these membrane invaginations, whether exocytic, endocytic, or simple membrane ruffles, must be determined with careful experiments (Watanabe et al., 2013a, 2013b, 2014). If any of these pits are clearly covered with electron-dense materials indicative of clathrin-coats (Heuser and Reese, 1973), they can be annotated as clathrin-coated pits (“7”). Note that the active zone membrane is drawn under the pit where the plasma membrane would have been before exocytosis, when prominent pits like the one in Figure 1B are present within the active zone. The Matlab routine distinguishes pits within the active zone (Figure 1B) from those outside (Figure 2E,F) by whether two ends of the segmented line for pits are within the traced active zone membrane. If both ends are within 5 nm of the active zone membrane, they are considered being within the active zone. Thus, these features must be traced carefully.

**Figure 2.**
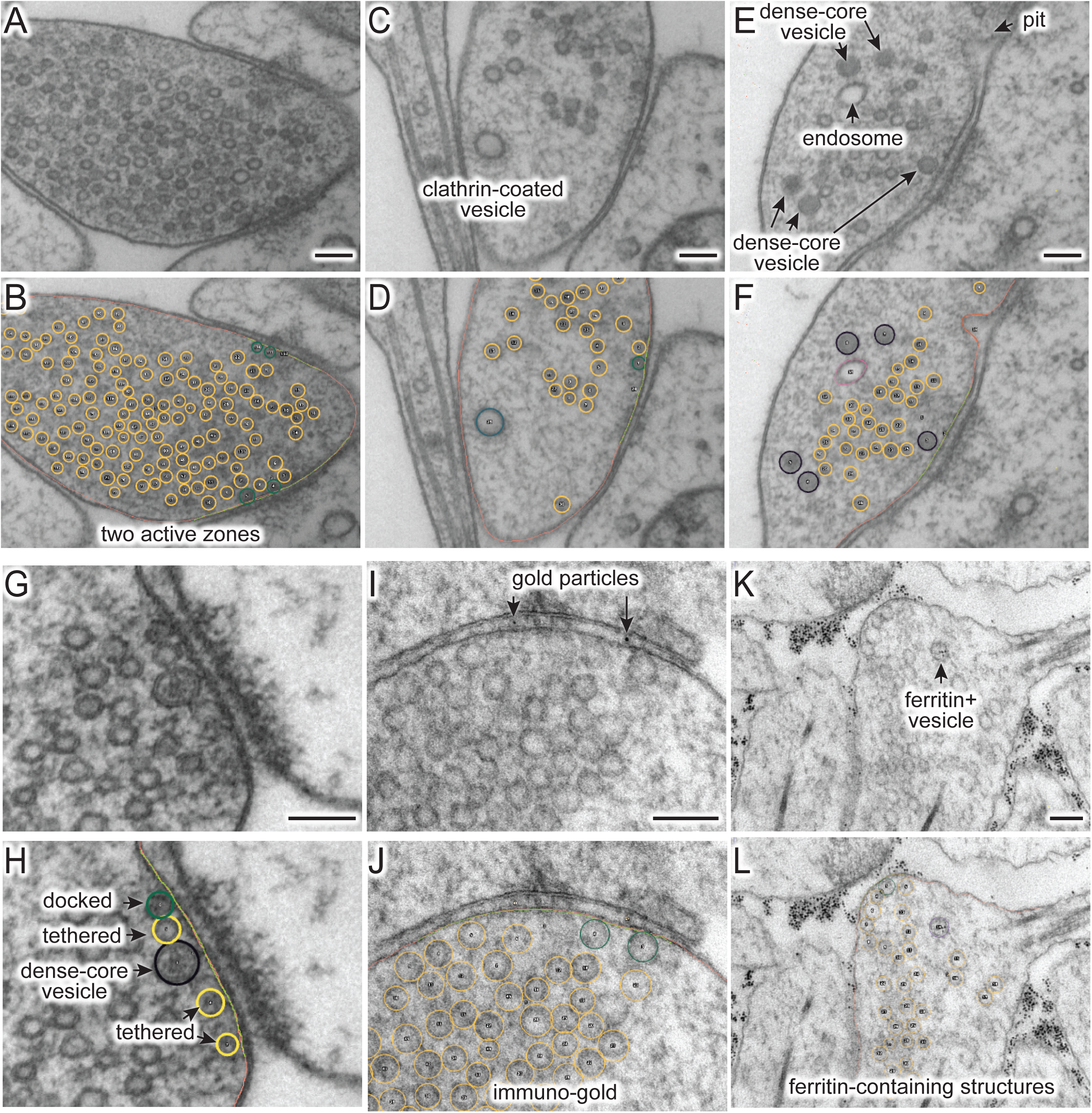
(A-L) Example micrographs (A, C, E, G, I, K) and their annotations (B, D, G, H, J, L), showing structures that can be traced using the ImageJ/Fiji macros (“synapsEM_analysis_macro.txt”). (A, B) Multiple active zones in a single synaptic profile. (C, D) Clathrin-coated vesicle. (E, F) Dense-core vesicles and endosomes. (G, H) Docked or tethered synaptic vesicles in active zones. (I, J) Gold particles. (K, L) Ferritin-containing vesicles. See Table 2 for the full list of structures and hot keys to enter into the ROI manager. Scale bars = 100 nm.

For vesicles, use the straight line tool to draw a line across the outer edges of a vesicle (Figure 1D). By doing so, the diameter of a vesicle as well as the x,y-coordinates at the center of the line are recorded in the ROI manager. A circle is drawn on the vesicle based on the diameter. Synaptic vesicles can be manually categorized (Figure 2G,H) into docked – no lighter pixels between vesicle membrane and plasma membrane (“3”), tethered when a vesicle is close but not docked and has visible tethers to the plasma membrane (“2”), and all other vesicles in the terminal (“1”; Figure 2G,H). If not categorized, docking will be determined by the overlap between the vesicle membrane and plasma membrane. However, tethered vesicles must be annotated by visual inspection, since they are defined as having a physical tether to the plasma membrane (Watanabe et al., 2013b). The same line tool is used to annotate other types of vesicles including clathrin-coated vesicles (“8”, Figure 2C,D), dense-core vesicles (“4” and “5” if docked, Figure 2E-H), and large vesicles (“6”, Figure 1B). Large vesicles are clear-core vesicles with a diameter of 60-100 nm that may be involved in exocytosis (Borges-Merjane et al., 2020; He et al., 2009; Maus et al., 2020), endocytosis (Kononenko et al., 2014; Watanabe et al., 2013a, 2013a, 2014), and cargo trafficking (Ou et al., 2010; Vukoja et al., 2018). If preferred, all vesicles can be annotated as synaptic vesicles using “1”.

Endosomes are also quite prevalent at presynaptic terminals. Although their identity is difficult to determine from single profiles, we define structures as endosomes if they are larger than 100 nm by visual inspection or have irregular shapes (Figure 2E,F) (Watanabe et al., 2014, 2018). To trace endosomes, the freehand selection tool (“F4”) is used to follow the contour of the putative endosomal membrane, and press “0” from the numeric keypad to add the coordinates (“n0” – hereafter, when a number is preceded by n, it will refer to the number key on the numeric keypad). Their areas are also recorded in the ROI manager. For multivesicular bodies (MVBs), press “m”. Currently, endosomes and MVBs are the only such structures traced in our study, but this analysis can be extended to other irregularly shaped structures like mitochondria and autophagosomes.

Ferritin or gold particles, as well as organelles that contain them can also be tracked as distinct structures (Figure 2I-L). Ferritin is typically used as a fluid phase marker to track recently endocytosed membranes (Watanabe et al., 2013b, 2014, 2018), while gold particles are often conjugated with antibodies or other moieties for affinity interaction to probe proteins of interest (Li et al., 2020). In addition to the plasma membrane and active zone membrane, pits with particles can be annotated using the freehand line tool and recorded with “n6” on the numeric keypad. For particle-containing vesicles, use the straight line tool, and add them to the ROI manager by pressing “n1” for any vesicles, “n2” for tethered vesicles, “n3” for docked vesicles, “n4” for large vesicles, and “n5” for clathrin-coated vesicles (Table 2). The particles themselves can be annotated using the same tool and hitting the “p” on the keyboard. Particle-positive endosomes and MVBs are traced using the freehand selection tool and recorded with “n8” and “n7”, respectively.

For other structures, the best practice is to annotate using hot keys of structures similar to the structures of interest. For example, mitochondria can be marked as “particle-positive MVBs (n7)” or “particle-positive endosomes (n8)” for the analysis purpose. Likewise, any vesicular structures can be annotated with particle-positive vesicles (n1-n5) since these structures are normally not annotated unless ferritin or something similar is used in the study. Be sure to take notes on what keys are used. As a cautionary note, the distance between membranes is calculated at the minimum, that is, from the outer leaflet of a vesicle to the inner leaflet of the plasma membrane or active zone membrane. Accordingly, it is highly recommended to use images acquired at a sufficient spatial resolution (i.e. less than 1 nm/pixel) and use a pen tablet to trace objects. Since electron micrographs cannot be acquired at the same settings, the contrast should be adjusted in each image to make the features-of-interest appear clearer for analysis.

### 3.4 Exporting as a text file

When segmentation is completed from an image, the annotated structures in the ROI manager can be exported as a text file. Press “e” on the keyboard (Figure 1B). This macro then generates a text file containing all the segmented structures in the order of the ROI manager list. The text file is named after the image and automatically saved in the folder where the image is. The Fiji screen advances to the next image, after the text file is saved. In the text file, the record of each structure is organized as follows: the tool used to annotate (by its Fiji tool number), the name of a structure, area or length of a structure if it is not a vesicle, x-coordinate(s), y-coordinate(s), and radius of a vesicle if it is a vesicle. These values are separated by a tab character (ASCII 09). The record in the ROI manager will be erased after the export is complete.

### 3.5 Importing a text file

After completing the analysis for all images, it is important to check whether all the essential components are segmented in every image and the text files are compatible with the Matlab codes. To check, run the “start_data_check” function in Matlab (Figure 1C). If any data are missing or more than one plasma membrane are annotated, this function returns the names of the files and the description of the problems associated with the files in the command window. To fix the problems or reevaluate the annotations, the records in a text file can be imported back to the correct image in a stack. On the image of interest, press “i” to import the text file. If there is a text file corresponding to this image in the folder, the segmentation will automatically pop up and the records in the ROI manager. If the list in the ROI manager is modified, press “e” again to export the modified list to the text file. Note that this process will overwrite the existing file.

### 3.6 Unblinding the text files

After ensuring that all images are annotated correctly, the resulting text files can be unblinded for further analysis by the Matlab scripts (Figure 1C). To unblind the shuffled text files, use the “unblind_me” script in the Matlab. This script prompts users to define the directory where the key is and where the text files are. After selecting the files, the text files will be renamed based on the names of the original images and copied into a new folder, called “unblinded”, nested within the folder with the shuffled text files. Further analysis should be performed with the unblinded text files. Both blinded images and text files are untouched during the process, and thus, the experimenters can be blinded for further analysis.

### 3.7 Running Matlab scripts

The Matlab scripts are designed to use the spatial coordinates and size information of annotated structures to calculate the distances between them, and extract the essential parameters for synapse morphometry. To start, select the directory where all the scripts are located, and then type in “start_analysis” in the command line. Since the data analysis for each sample must be run separately, we typically define the name of the sample at this stage (i.e. control_1 = start_analysis;). The program prompts users to input the pixel size for the images (nm/pixel) and the size of bin (i.e. 50 nm), which is used for plotting the distribution data such as locations of vesicles relative to the active zone. Then, a user interface pops up, first asking to choose the directory where the text files are and then to select all the text files. The files are then loaded into the Matlab for processing. The scripts are designed to perform the following three calculations: size, number, and distribution, of structures.

First, the sizes of structures are calculated based on the pixel size, and the mean and median size of vesicles and endosomes are determined from the sample. These numbers are available as an average in a single image or an average for the sample. For pits, the diameter is calculated at the full-width half-maximum. In addition, the scripts also calculate the depth (height), the width at the base, and the surface area of the pits. These data are available as a number array in the final dataset. Second, the total numbers of structures are calculated from each profile and then their mean and median are determined for the sample. The total numbers are additionally sorted based on their locations relative to the active zone and plasma membrane. For example, if a vesicle is within 30 nm of the plasma membrane and also within 30 nm of the active zone, this vesicle is counted towards the vesicle associated with the active zone. If a vesicle is within 30 nm of the plasma membrane but not associated with the active zone membrane (> 30 nm), this vesicle will be categorized as a vesicle in the periactive zone. If neither condition is met, the object is considered cytoplasmic. The distinction between vesicles above the active zone versus periactive zone is somewhat arbitrary but useful for detecting certain vesicle pools. For example, synaptic vesicles within 30 nm of the active zone membrane (about two rows of vesicles) are thought to contribute to the readily releasable pool (Schikorski and Stevens, 2001), and their numbers are often reported (Hammarlund et al., 2007; Richmond et al., 2001). Vesicle or pits in the periactive zone reflect endocytic events since they correlate with the internalization of fluid phase markers and are typically observed hundreds of milliseconds after an action potential (Watanabe et al., 2013b, 2014, 2018). In contrast, pits within the active zone represent fusing vesicles since they appear a few milliseconds after an action potential (Kusick et al., 2018; Watanabe et al., 2013a, 2013b). The distance threshold in nanometers can be moved by changing the number in line number 65 in the source code (vesicle_count.m) from 30 to the desired number. After the analysis, the number data are available in the “vesicle_number” table as a number array. The key to interpret the array is listed in Table 4.

Third, they calculate the minimal distance from each structure to the plasma membrane and active zone membrane and determine the distribution of each structure relative to these membranes. For a vesicle, the distance from the center to every point on the plasma membrane, active zone, and if annotated, dense projection is calculated, and the radius of the vesicle is subtracted such that the distance is determined from the outer edge of the vesicle to the membrane. Then, the minimal distance is reported as the final distance. If the distance is calculated to be 0 nm from the plasma membrane, the vesicles are considered docked, and they will be categorized into the docked pool for the number calculations. For endosomes, distances are calculated from every point of their membrane to every point on the plasma membrane, active zone, and dense projection, and their minimal distance is reported. The distances can be plotted as continuous frequency distribution with no binning, if enough data are collected. However, the distribution of structures is typically determined by calculating the number of the structures at certain distances away from the active zone membrane or if annotated, the dense projection based on the bin the user specifies (i.e. 50 nm). The resulting tables show their average number and normalized abundance at each bin (Table 3).

The output of the Matlab scripts appear in the workspace as a structure array and can be compared between different conditions or samples (these can be control vs mutants, or glutamatergic vs GABAergic neurons). To compare, all other samples in a single experiment should be processed by the same procedure (i.e. mutant_1 = start_analysis; in the command line). After all the samples are processed, the workspace should be saved in .mat format.

For plotting the data and statistical analysis, we export the data to Prism. Several optional scripts are available to re-organize the data for exporting. Please refer to the Supplementary Information. Step-by-step protocols are also available in the Supplementary Information.

## Results

To validate program scripts, the computed data were compared to data segmented manually. Specifically, we segmented 10 images using the procedures described above and calculated distances from ∼25-30 vesicles to the active zone membrane using the Matlab scripts. Then, we manually measured the distances from those vesicles to the nearest active zone membrane based on visual inspection. We repeated the measurements three times to estimate errors caused by manual measurement. We then plotted the disparities between distances determined by the different methods. On average, the difference between the calculated and measured distances was 1.4 nm (Figure 3B; median and 95% CI). This number is similar to the error made by repeating the manual measurements three times on the same set of images (1.3 nm median, Figure 3B). Thus, the calculations based on the x,y-coordinates of structures are valid and produce data as accurate as manual measurements.

**Figure 3.**
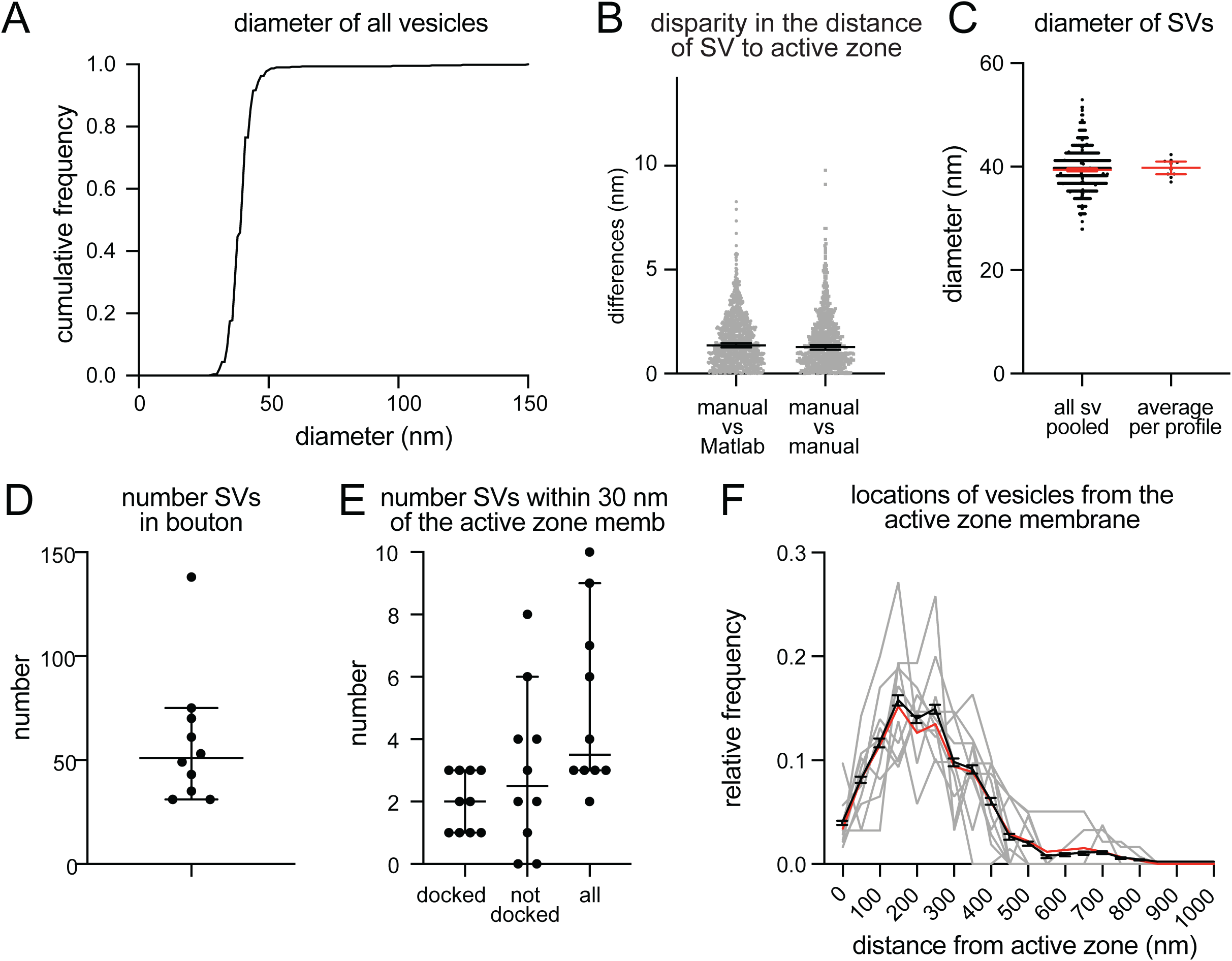
Example plots that can be readily generated with the synapsEM. (A) A cumulative plot showing diameter of all vesicles annotated from 10 sample images used in this study. A total of 590 vesicles are annotated. (B) A scatter plot showing the disparity in distances of vesicles to the active zone membrane between the Matlab calculated and manually measured (left) or among three manually measured (right). Each dot = one measurement. The medians are 1.4 nm and 1.3 nm respectively (p = 0.8, unpaired t-test). (C) A plot showing the diameter of synaptic vesicles, averaged from all vesicles pooled (left, 39.6 ± 0.2 nm, mean ± SEM) and means of each profile (right, 40.1 ± 0.6 nm, mean ± SEM). (D) A plot showing the number of vesicles in the terminal. Each dot = the number in each profile. (E) A plot showing the number of vesicles, docked or tethered at the active zone or all vesicles within 30 nm of the active zone membrane. Each dot = the number in each profile. (F). A plot showing the distribution of synaptic vesicles relative to the active zone. Gray lines = the normalized distribution from each profile. Black line = average from 10 profiles. Red line = the normalized abundance from the pooled data.

Based on the data from these 10 images, we determined key synaptic parameters of synapses from cultured mouse hippocampal neurons (14 days *in vitro*). The diameter of synaptic vesicles was 39.6 ± 0.2 nm (mean ± SEM), when all vesicles in the set were pooled, and 40.1 ± 0.6 nm (mean ± SEM), when the numbers were first averaged per profile and then the mean of means was determined from the entire dataset (Figure 3C). The mean of means would better represent the population average. The average number of synaptic vesicles per profile was 51 (Figure 3D, median and 95% CI shown). About 4 vesicles were found within 30 nm of the active zone membrane; of these, ∼2 vesicles were docked on average (Figure 3E). It is sometimes useful to normalize vesicle numbers and docking to the size of active zone. To accommodate this calculation, the length of active zones is accessible in the output (Table 4), and average is also determined (median = 385 nm). Although n is small, these numbers are surprisingly close to the numbers we have obtained from thousands of images across many experiments (Watanabe et al., 2013, Watanabe et al., 2014, Kusick et al., 2018, Li et al., 2020). The distribution of synaptic vesicles relative to the active zone can be mapped from each synaptic profile (gray lines), averaged numbers per profile (black line, mean ± SEM), and the data all combined (Figure 3F, red line). Thus, typical synaptic parameters can be measured and plotted using SynapsEM.

SynapsEM can be used to segment data from other model systems. We performed the same analysis using serial sections of *C. elegans* neuromuscular junctions (Figure 4, n = 5 reconstructed synapses). In these reconstructions, the numbers can be calculated per synaptic profile containing a dense projection or per fully reconstructed synapse (end-to-end of a synaptic bouton defined by the presence of synaptic vesicles). The average number of synaptic vesicles per profile and per reconstructed synapse were 35.5 and 394, respectively (Figure 4 A-C, E). Of the ∼5 vesicles within 30 nm of the active zone membrane in single profiles, an average of 3 are docked per synaptic profile (Figure 4D), thus, 8.5% of the total vesicle pool are docked in the active zone. Similarly, 9% of the total vesicles in the reconstructed boutons were docked (Figure 4F). Thus, the docked pool can be estimated from the synaptic profile data, as has been done in previous publications (Hammarlund et al., 2007; Watanabe et al., 2013a). Since dense projections are apparent at these synapses (Figure 4A), locations of synaptic vesicles and docked vesicles can be mapped relative to the dense projection (Figure 4G, H). The median distance for all vesicles was 140 nm, while that for docked vesicles was 67 nm, suggesting that vesicles tend to dock near the dense projections, where voltage-gated calcium channels are harbored (Gracheva et al., 2008). Thus, SynapsEM works with both 2D and 3D datasets from multiple model systems.

**Figure 4.**
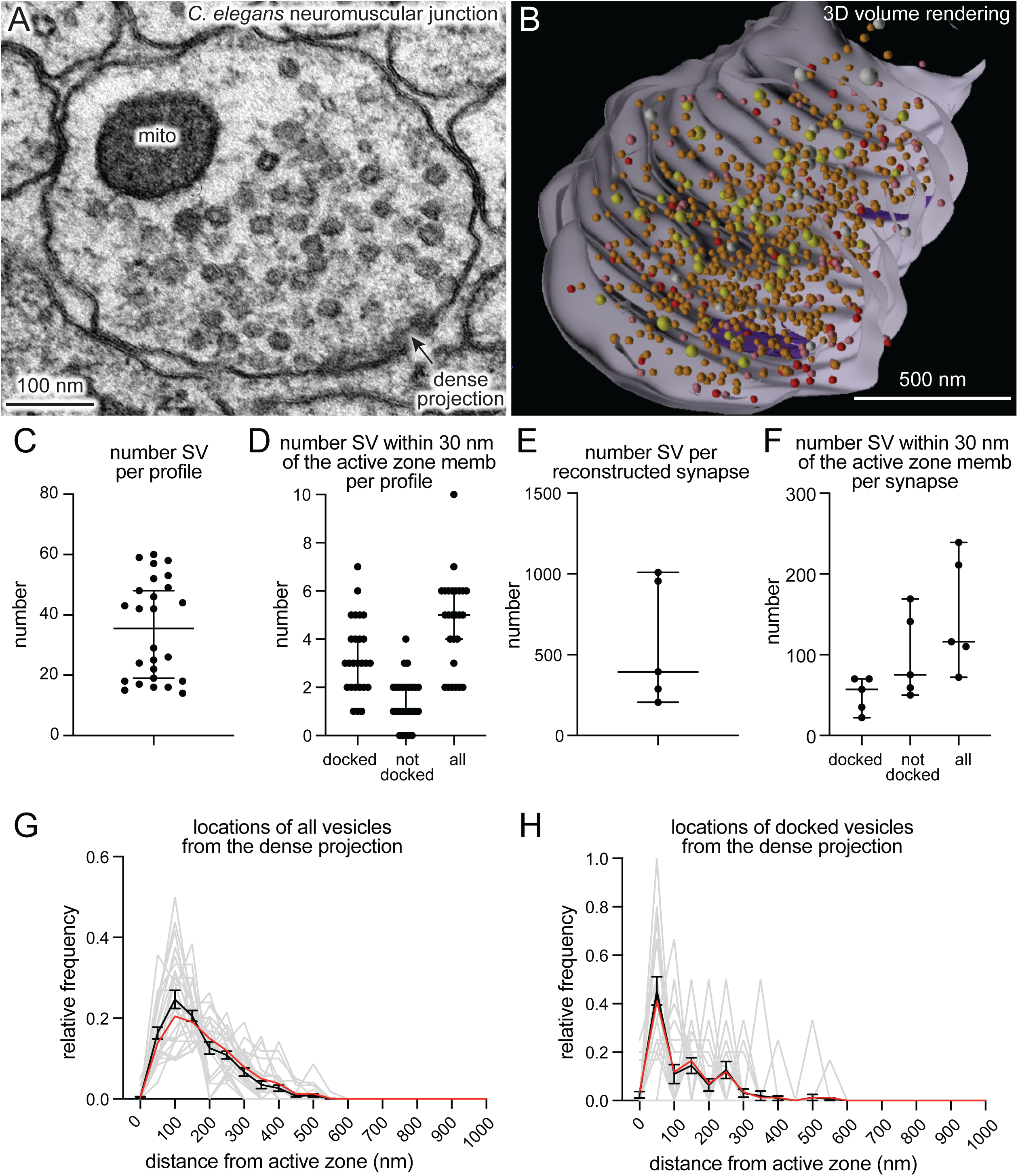
SynapsEM works with other model systems and with 3D reconstruction programs. (A) An example micrograph of a *C. elegans* neuromuscular junction. (B) A snap-shot of the reconstructed synapse using Maya, based on the x,y-coordinates collected from each profile in this study. Red = docked synaptic vesicles; Orange = synaptic vesicles; yellow = dense-core vesicles; white = large vesicles; purple = dense projections. (C) A plot showing the number of vesicles in single synaptic profiles. Each dot = the number in a profile. (D) A plot showing the number of vesicles within 30 nm of the active zone membrane per profile, and the number of those vesicles docked or tethered at the active zone. Each dot = the number in each profile. (E) The total number of synaptic vesicles from synapses fully reconstructed from serial electron micrographs. (F) the number of vesicles within 30 nm of the active zone membrane per profile, and the number of those vesicles docked or tethered at the active zone from fully reconstructed synapses. (G) A plot showing the distribution of synaptic vesicles relative to the dense projection. (H) A plot showing the distribution of docked synaptic vesicles relative to the dense projection. In G and H, gray lines = the normalized distribution from each profile. Black line = average from 26 profiles. Red line = the normalized abundance from the pooled data.

## Discussion

Ultrastructural analysis of synapses has been performed by many labs over the years. Excellent programs, for example IMOD (Kremer et al., 1996), TrakEM2 (Cardona et al., 2012) and Reconstruct (Fiala, 2005; SynapseWeb, Kristen Harris), provide visualization software for data acquired from either serial sections or tomograms. SynapseEM is a Fiji plug-in designed to quantify morphometric data from electron micrographs. IMOD, TrakEM2, and Reconstruct can all perform similar measurements but focus more on 3-D data, with in-depth features for handling and rendering serial images or tomograms not included in SynapsEM. TrakEM2 in particular offers similar quantification procedures; SynapsEM’s benefit is an all-in-one package, from scrambling raw images all the way to outputting final data tables, that is fast and easy to use even for those completely unfamiliar with Fiji or Matlab, but also easily modified. These procedures have been used by everyone from novices to experts to quantitate tens of thousands of 2-D images and over a thousand 3-D reconstructions (Kusick et al., 2018; Li et al., 2020; Watanabe et al., 2013b, 2014, 2018). Although SynapsEM handles serial-section data, it is difficult to annotate structures like endosomes and multivesicular bodies that span across multiple sections, since images are randomized. Thus, for serial-section data, careful re-evaluation is necessary after unblinding.

Several features streamline and standardize characterization of synaptic features. First, this approach allows multiple experimenters to assess the validity of annotations, reducing potential errors in the analysis. Second, automated shuffling of images from different conditions reduces potential bias in the analysis. Third, additional parameters can be extracted from the dataset *post hoc*, since the positions of structures are all recorded. For example, the locations of pits relative to the center of an active zone can be calculated based on their coordinates (Kusick et al., 2018; Li et al., 2020; also see Supplementary Information). Fourth, this approach can also be applied to serial-sections to calculate distances in three-dimensions (Kusick et al., 2018), similar to IMOD (Kremer et al., 1996) and Reconstruct (Fiala, 2005; SynapseWeb, Kristen Harris). Fifth, the 3D dataset can be rendered into a segmented volume in Maya based on the x,y-coordinates of the structures and their sizes (Figure 4B; see Supplementary Information for the procedure). Thus, SynapsEM is a versatile approach for the morphometry of synapses. For truly universal and automated analysis, the implementation of the machine-learning based algorithms into the SynapsEM platform is awaited. It is hoped that SynapsEM will promote data sharing and consistent morphometry of synaptic ultrastructure among labs.

## 7 Author Contributions

S.W., M.W.D., and E.M.J. conceived, designed the experiments, and wrote the manuscript. G.F.K. and S.W. wrote the protocols. M.W.D. wrote the original macro for ImageJ/Fiji. S.W. wrote the Matlab scripts. J.I. wrote the Maya code for volume rendering.

## 9 Acknowledgments

We thank H. James de St. Germain for training S.W. on Matlab coding and Marc Hammarlund for discussion. We are also indebted to Edward Hujber, Thien Vu, Morven Chien, and Stephen Alexander Lee for additional MatLab routines.

## 11 Data Availability Statement

The datasets analyzed for this study can be found in the https://figshare.com/account/home#/projects/85250. The scripts are available from https://github.com/shigekiwatanabe/SynapsEM.

