## supplementary information for "SynapsEM: computer-assisted synapse morphometry"

Step-by-step protocols for running SynapsEM are described below. The **bald** font indicates the actions you need to take. More descriptions are in regular fonts and also found in the Methods section of the manuscript. The texts in red are cautionary notes.

Randomizing images

**1. Getting started with Matlab**:

**1.1 Open Matlab.**

-At the center is the Command Window, where code is run. Scripts are called by typing in their name (for example, for "randomize.m", type "randomize") and pressing enter/return. Scripts will sometimes ask for input while they are running (for example, a yes/no question: "do you want to duplicate the images y/n", or asking for a number input "what is the pixel size?"). Simply type the response into the Command Window and press enter/return to continue running the code.

-At the right is the Workspace. This is where Matlab stores "variables": for SynapsEM, this is where your data will be stored. The "Save Workspace" button in the "Home" panel of the top toolbar will save all the current variables as a .mat file.

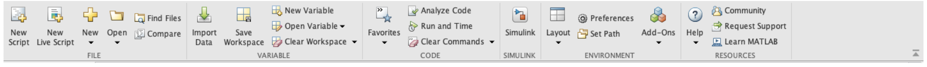


-At the left is the Current Folder. This is the directory where the Command Window can call scripts (.m files) from. Before you get started, you will need to set the directory to where you have SynapsEM code stored.

-When you open up something in the Current Folder (to edit/look at scripts) or the Workspace (to look at data), the Command Window will move to the bottom of the screen.

**2. Setting the Current Folder to where you have the code stored:**

**2.1** At the top of the screen, below the toolbar, you will see the file path to where the Current Folder is currently set to. **Click through the file path until it ends with the folder where you have all SynapsEM code stored**. For example:

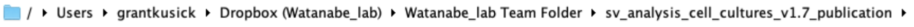


-The codes should pop up under Current Folder. Your screen should look like this:

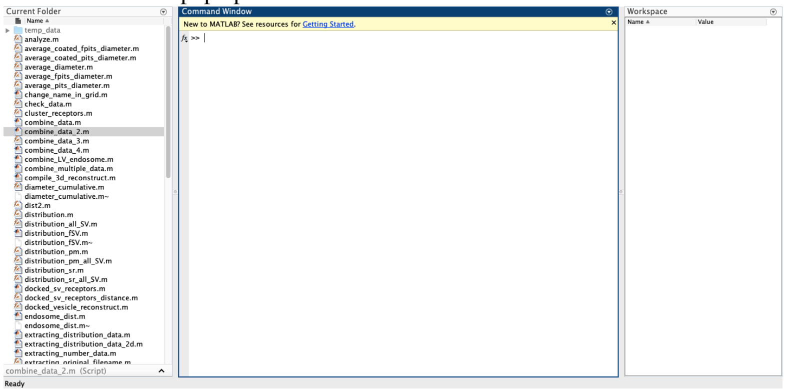

-Matlab will remember your Current Folder after closing, so this step doesn't need to be repeated each time it's opened.
-Scripts can be inspected or edited by double clicking on the .m files. This will open a "script editor" panel. SynapsEM scripts are heavily commented (in green text), so inspecting these files can be helpful if you are troubleshooting or simply want to know more about how the scripts work.

**3. Randomizing images:**
**3.1 Move all your image files for a single experiment into a single folder, or duplicates of these files.**
**3.2 In Matlab, run the "randomize" script*.**
**3.3 Select the folder that contains the images**
**3.4 Select all the images that you want to randomize.**
-The code will ask whether you want to duplicate the images. Answering yes will make a copy of the files and randomize them. Answering no will randomize the names of the files you selected
Warning: if you answer "no" the code will overwrite the original file names of the files you selected. If you have not already made copies of the files, make sure to type "yes".

```
>> randomize
Select a folder containing tif files
do you want to duplicate the images? (yes/no)
```

-The code will create a new folder called "randomized" within the folder you selected. This should contain the same images you selected but with scrambled file names.

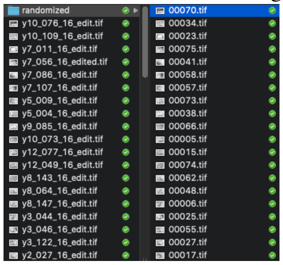


-Important: the randomized folder should also contain a Matlab file called "key". Do not lose this file: you will need it to unscramble your text files later.

**4. Getting started with Fiji:**
**4.1 Open Fiji**
**4.2 Select "Install"**

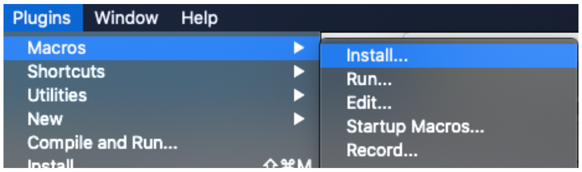

4.3 **Install the file "synapsEM_analysis_macro.txt"**
-This process has to be done each time you boot up Fiji. To avoid this, you can copy the text in synapsEM_analysis_macro.txt into the text file "StartupMacros.fiji.ijm". This is the list of macros that Fiji calls by default when opened.

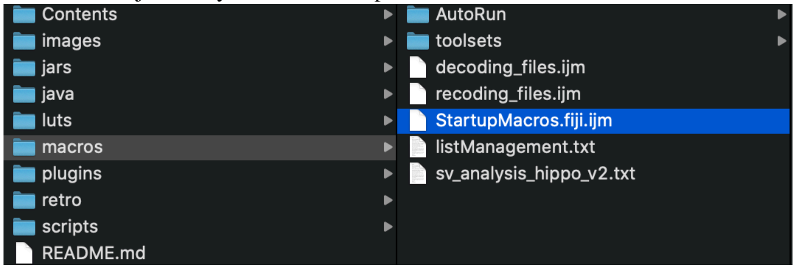


-Now when you go to "Macros" it should look like this:

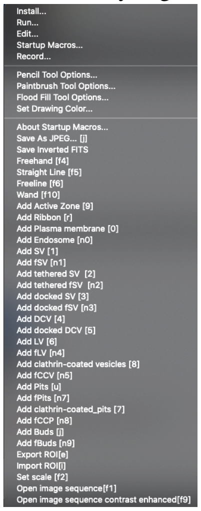


This is a list of the macros and the keyboard shortcuts used for each. "n" indicates the number pad. For example pressing "0" on the top number row of your keyboard will execute the add plasma membrane macro, whereas pressing 9 on the number pad will execute the add fBuds macro. Refer **Table 2** for a complete set of macros.

4.4 **Open a stack of your scrambled images by pressing F1.** Important: you cannot simply drag and drop images into Fiji, you must use the F1 macro. Note: if you are not sure if you used F1 to open images, press F2 to set the scale before stating the annotation.

**5. Segmenting images**

Segment features as described in the Methods section and corresponding figures.

**5.1 Press F6 to activate the freehand line tool.**

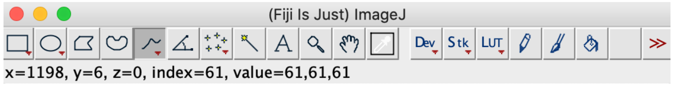


**5.2 Trace the contour of the plasma membrane and press "0" on the keyboard.** Important: you can only annotate one plasma membrane per image. Make sure "show all" and "labels" are selected in the ROI manager so you know which structures are already segmented.

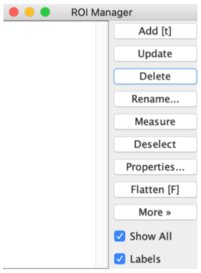


**5.3 Trace the contour of the active zone membrane and press "9" on the keyboard.** You can select multiple active zones if necessary.

**5.4 Press F5 to activate the straight line tool**.

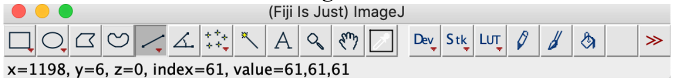


**5.5 Draw a line across a vesicle (outer edge to outer edge) and press appropriate numbers on the keyboard (see Table 2).** For examples, press "1" for a synaptic vesicle and "3" for docked synaptic vesicles. **Repeat this procedure until all vesicular structures are annotated.**

**5.6 Press F4 to activate the freehand selection tool.**

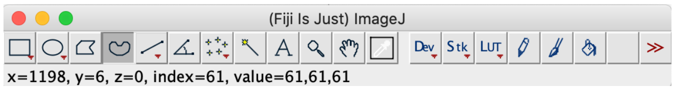


**5.7 Trace the contour of an endosome or multivesicular body and press "n0" on the numerical keypad for an endosome and "m" for multivesicular body.**

The order of 5.1-5.6 is not important, but for the Matlab program, exactly one plasma membrane and at least one active zone must be annotated.

### 6. Exporting text files

**6.1 Press "e" on the keyboard.** The macro generates a text file containing all the segmented structures in the order of the ROI manager list. The text file is named after the image and automatically saved in the folder where the image is. The Fiji screen advances to the next image, and the text file is saved. In the text file, the record of each structure is organized as follows: the tool used to annotate (by its Fiji tool number), the name of a structure, area or length of a structure if it is not a vesicle, x-coordinate(s), y-coordinate(s), and radius of a vesicle if it is a vesicle. These values are separated by a tab character (ASCII 09). The record in the ROI manager will be erased after the export is complete.

**Repeat 5-6 until all images are annotated.**

**7. Checking data**
Before running the Matlab analysis script, you need to check that no errors are in the data that would cause the code to fail to run (missing active zone, multiple plasma membranes, etc.).
**7.1 Run the script 'start_data_check'.**
**7.2 Select the folder with the files and then the files themselves.**
-The script will scan through all the files to see if there are any issues that would cause the analysis code to fail: if the analysis code (described below) has an error caused by the text file, it will not tell you what caused it or which text file had the issue, but start_data_check will. Below is an example of the script running and detecting errors, indicating the type of problem to be fixed and which files have the problems.

```
Is this for ribbon synapses? (yes/no) n
Choose the directory where analysis data is loated
/Users/grantkusick/Dropbox (Watanabe_lab)/Watanabe_lab Team Folder/Working Data/Grant/EGTA_docking_2_pix_scrmbl/3.584264137782
too many plasma membrane data
/Users/grantkusick/Dropbox (Watanabe_lab)/Watanabe_lab Team Folder/Working Data/Grant/EGTA_docking_2_pix_scrmbl/169.4057952435
plasma membrane missing
```

-Fix any errors in the text files as needed (see section **8**).

**8. Importing text files (re-annotating images/correcting analysis)**
**8.1 Press 'i'.** Provided the text files and images are still in the same folder and still have the same name, the segmentation will appear on the image. You can now edit the segmentation as desired. Re-checking segmentation, ideally by another member of the lab, is important to ensure accuracy. You can also use this procedure to annotate features that you ignored in your first round of segmentation.
-To view segmentation that has already been done on an image, you must first open an image stack using the f1 macro.
-Export by pressing 'e'. Important: this will automatically overwrite the original text file. If you wish to keep the original files, make copies of them and move them to another folder before re-annotating images.

**Repeat 7 to make sure all the text files are compatible.**

**9. Unscrambling text files**
**9.1 Open Matlab and call up the SynapsEM scripts as described in section 2**.
**9.2 Run the script 'unblind_me' by typing in the command window.**
**9.3 Select the folder containing the text files and decoding key.**
**9.4 Select the text files.**
-The text files will be renamed based on the names of the original images and copied into a new folder, called "unblinded", nested within the folder with the shuffled text files. Further analysis should be performed with the unblinded text files.

**10. Running analysis scripts**
**10.1 Run 'start_analysis'.** This generates all of the data you will need for most experiments, and these numbers serve as the basis for any other analysis scripts you might want to run. When prompted, choose all the text files corresponding to a single sample: for each run MATLAB generates an object in the workspace that contains all the data for that sample.  You will be asked for 1) the pixel size in nm (all data in MATLAB will be in nm) and 2) the bin size, which for

data relating to the distance of objects (SVs, LVs, etc.) to the active zone/plasma membrane will determine how these distances are binned. (in 2 nm increments, in 20 nm increments, etc).
- Name the object after that sample, either by changing it after it's generated (control click on the folder in the Variables window and select rename) or by indicating it in running the code, for example "mutant_1= start_analysis". Important: if you do not specify a name, the data set will have the name "ans" by default. If you run the script again for a new set of txt files, it will overwrite the first data set. Therefore, make sure to name the data set before adding the next data set in.

**10.2 Repeat for each sample in the experiment, making sure the name of each data set is correct.**

**10.3 Save the data by using the "Save Workspace button", in the Home window of the top toolbar.**

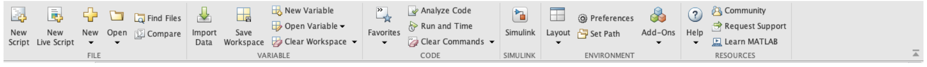


-This will save a .mat file, which is all of these data sets ("Variables") saved together. Whenever opening a .mat file, you will be asked which of the variables you want to load in; all will be selected by default. The easiest way to keep things organized is to keep one .mat file for each experiment.

-Your Workspace should now look something like this:

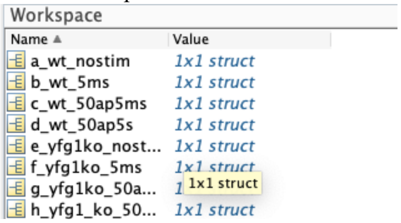


-Once saved, you can remove folders from the Variables window to reduce clutter as desired, or clear the Variables window by inputting the command "clear all".

**11. Examining data**

**11.1 Double click on any of the folders in the Variables window to open up that data set.**

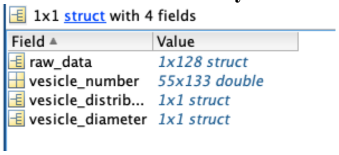


-Refer to Table 4 for how the features are arranged in the vesicle_number data table.
-Refer to Table 1 for how features are arranged in the vesicle_distribution tables.
-Move data into your statistical analysis/data visualization platform of choice
-Enjoy the thrill of discovery!

**12. Specialized scripts: making data into tables for export**

You can export count data manually out of the spreadsheets, but this can be streamlined using the ‘extracting_number_data’ script.

**12.1 Once you run through all your data, change the names in the workspace so that they are in the order you want to have in your final graph.** For example:
control no stim
control 100 ms
control 1s
control 10 s
kd no stim
kd 100 ms
kd 1s
kd 10s

-The easiest way to make it in this order is to add alphabet in the beginning.

a_control no stim
b_control 100 ms
c_control 1s
d_control 10 s
e_kd no stim
f_kd 100 ms
g_kd 1s
d_kd 10s

**12.2 Once you do so, save the workspace (see 10.3).**
**12.3 Then in the command window, type in extracting_number_data.** It will prompt you to answer 2 questions. Answer yes to the first question if you need usual numbers like docked synaptic vesicles, pits, large vesicles, endosomes, etc. Answer yes to the second question if you want to get ferritin-containing structures, too.

If you would like something special like clathrin-coated pits, say no to both questions. It will ask you to type in the row number you want to get the values from. In the case of clathrin-coated pits, this would be 46.

It should give you sample_out and table in the workspace. If you answered yes to either question, go to sample_out. If not, go to table. The first row is the total number of sections you analyzed for the particular dataset. The columns are organized according to how the names were organized in the workspace (see above, i.e. 1st column, control no stimulation, 2nd, control 100 ms, etc). You can copy the whole table except for the first row.

*The only problem is that the matlab will fill in empty slots with 0s, so if the total number of images analyzed is not uniform across, you will need to remove those extra 0s based on the total number you analyzed for each group. For example, out of the 8 samples mentioned above, if control no stim had 90 images, but others had 100, the Matlab will add ten zeros to the control no stim. Remove those 0s after you copy the data out.

**12.4. To extract the size data, type in "extracting_size_data" in the command window and select the directory and saved dataset (.mat).** Sample_out shows up in the workspace. Double-click on sample_out and navigate through the variables to access the data. For example, for synaptic vesicle diameter, click on "SV". Two tables are available: "pooled_raw_number" and "mean_profile". The pooled_raw_number pools diameter of all vesicles from every profile in the sample, which is used to generate a plot in **Figure 3C**. Each column represents every vesicle in this case. The mean_profile lists the mean of synaptic vesicles from every synaptic profile. Each column represents the mean diameter of all synaptic vesicles in a profile. The first row is the total number of sections you analyzed for the particular dataset in both cases. The columns are organized according to how the names were organized in the workspace (see above, i.e. 1st column, control no stimulation, 2nd, control 100 ms, etc). You can copy the whole table except for the first row. Again the 0s will need to be eliminated (see 12.3). The diameter data for vesicles are also available as cumulative frequency, as shown in **Figure 3A**. In this table, The columns are also organized according to how the names were organized in the workspace (see above, i.e. 1st column, control no stimulation, 2nd, control 100 ms, etc). Each row is 1-nm bin.

**12.5. To extract the distribution data, type in "extracting_sv_distribution_from_pm_data", or "extracting_sv_distribution_from_dp_data", or "extracting_sv_distribution_from_az_memb_data".** As the names of the scripts suggest, these would summarize the distribution of vesicles relative to plasma membrane, dense projection, or active zone membrane. Two tables are available in each case: "average" and "norm_abundance". As the names suggest, average shows the mean number of vesicles at each bin. The binning is shown in the column 1, and the numbers from each sample starts from the second column in the same order as how the names were organized in the workspace (see above, i.e. 1st column, control no stimulation, 2nd, control 100 ms, etc). The norm_abundance showns the normalized abundance at each distance instead. These data can be simply copied and pasted into x,y spreadsheet of the Prism with the first column in the x column. Since all the samples are processed with the same bin size, there will be no 0s at the end to remove. Note that the last row is any vesicle over 1000 nm from the plasma membrane, dense projection, or active zone.

**13. Specialized scripts: determining the location of features within the active zone/postsynaptic density**

Are you interested in whether your gold-labelled protein of interest is concentrated in the center of the post-synaptic density, or distributed throughout? Are you interested in where vesicle fusions are located within the active zone? These codes output count data on these features, either for 2-D or 3-D serial-section analysis and, more importantly, tell you where each feature is relative to the center of the active zone/PSD, and also calculate pairwise distances between your features of interest See Li et al., 2020 and Kusick et al., 2020 for example applications of these scripts.

**13.1 Make sure the data you are interested in have already had "start_analysis" run, saved as a .mat file (i.e. you've gone through part 10 of the main protocol).** Make sure there are no other variables in the Workspace, as these scripts by default will run on every variable in the Workspace, and if any are not in the expected format, it will trigger an error and the script will fail.

When running the start_xxx_reconstruct codes, make sure you don't have any important unsaved variables in your Workspace. If this code experiences any errors, it will wipe everything currently in the Workspace

**13.2 Run the start_xxx_reconstruct code (where xxx are either pits or docked_sv)**. There are various scripts that determine the locations of different objects within the active zone/post-synaptic density, which all function basically the same way. For this example, we'll use **"start_pit_reconstruct"**.

**13.3 Select the folder and then the .mat file you are analyzing.** You will be asked if this is for 3D reconstructions. See part 14 for info on dealing with serial-section data.

**13.4 Input pixel size.**

**13.5 Input the thickness of your sections.** This is only used if using 3D data, but it will be asked regardless.

**13.6 Input pit cutoff height.** This will be asked regardless of if you are looking at pits or not. If looking at pits, this will simply sort the pits (indicated by a 1 or 0 in the data table) by height/depth (ie. deep vs shallow), it will still include data on all pits in the active zone regardless of their depth/height.

The code will run through, in this case calculating data for each pit in the active zone. If the code runs properly, you should see "Sample in" and "Sample out" variables in your workspace.

**13.7 Open "Sample Out"**

In "xxx_Data" you will find a data table (ignore rows that are only 0s). Each row is a single active zone/synapseThe columns are as follows:

1.Distance to the edge of the active zone/PSD

2. Distance to the center of the active zone/PSD
3. *Normalized distance to the center. This is the value you are most likely interested in*
4. Pit height or vesicle radius
5. the side index (which side of the active zone pits or docked vesicles are)

**Commented [GK1]:** Forgot what these are

If there are another 5 columns in the same row, this is another pit in the same active zone.

**13.8 "Distance matrix" data table:** this contains all pairwise distances between pits/docked synaptic vesicles, etc. that are in the same active zone. These values are duplicated, mirrored along a diagonal. Simply copy out data from one side of the diagonal.

**14. Specialized scripts: Serial-sectioning analysis**

**14.1 Run "start_analysis" for each synapse (stack of images from a single reconstruction) from a particular sample/condition, then save the workspace.** You will have a workspace populated by all the different serial section stacks, each as an individual variable (data set), each named something like synapse 1, synapse 2, etc

**14.2 If interested in data from start_analysis: Run the extracting_xxx_data code of your choice, depending on what you are interested in.** You can also go through each synapse and pull out numbers manually, but with many serial section stacks this is tedious.

**14.3 If interested in reconstruct (location within active zone) data:**

**Run the start_xxx_reconstruct code of your choice, depending on what you are interested in**. The reconstruct codes can give you data on locations of pits, docked vesicles, etc. from serial section stacks. In fact, these codes were originally written to calculate positions of these objects within the active zone in 3D.

Do the same as in **13**, but when asked "is this for 3D reconstruction?" answer yes.
**14.4 Delete the "Sample In" variable and save the workspace.** This will allow the compile_3d_reconstruct code in the next part to run: you must have not variables in your workspace other than what you get from "Sample out". Going through each synapse individually is cumbersome, but the next step will compile them into a single data set.
**14. Compile the synapses from one sample into a single data set with "compile_3d_reconstruct". Simply run compile_3d_reconstruct and choose the .mat file that you saved in 14.4.** This will pool the pit/vesicle counts from all the synapses from a sample, as well as their locations, into a single data set.

**15. Specialized scripts: Aligning pits/docked synaptic vesicles with receptors.**
If you marked receptors with gold particles (Li et al., 2020), you can align pits or docked synaptic vesicles to those gold particles, found in the synaptic cleft.
**15.1 "start_pits_reconstruct" as instructed above (13.2-13.6). Delete the sample_in and rename sample_out as "pits" or "docked_sv". Save the workspace.**
**15.2 Run "start_docked_sv_reconstruct" as instructed above (13.2-13.6). Delete the sample_in and rename sample_out as "docked_sv". Save the workspace.**
**15.3 Run "start_particle_recustruct as instructed above (13.2-13.6). Delete the sample_in and rename sample_out as "receptor". Save the workspace.**
**15.4 Load all three .mat files into the workspace so that pits, docked_sv, and receptor are available in the workspace. Save the workspace.**
**15.5 Run "start_alignment_analysis". Follow the instructions in the command window.** The data will be stored in the sample_out. Click on either "xxx_pits.PitDist" or "xxx_docked_sv.SVDist". The table lists distances from every pit/docked synaptic vesicles to every gold particles (if three gold particles are in the synaptic cleft and 4 docked vesicles are found, a total of 12 distances will be listed).

**16. Specialized scripts: Volume rendering.** For volume rendering of the 3D reconstruction data as shown in Figure 4B, the location data need to be exported back to the text files. For Maya, the same structures must be in the same text file. To do so, **type "text_convert_maya" in the command window. When prompted, type in the thickness of sections (i.e. 40 nm).** This program will generate new folders with the name of the samples in the same directory where the .mat file is. These text files then can be imported into Maya for volume rendering.

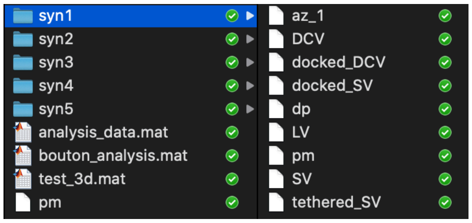


**17. Specialized protocols: Maya 3D modeling and rendering**.
**17.1 Modeling cells**: In Maya, closed curved are generated for each Z section. This is done by reading through the text file and running a "curve" command on x y points that share the same z

coordinate. To create a lofted surface, it is necessary for all of the curves to share the same number of vertices and to have their "seams" aligned. To accomplish this, the curves are selected and retopologized using the "rebuild" command with a specified number of control vertices (for example, 20), and their seams moved to face the same direction using the "move seam" command. The curves are then selected and used to create a polygonal mesh using the "loft" command.

**17.2 Modeling vesicles**: In Maya, a sphere is generated using the "sphere" command and moved to the x y z coordinates specified by the data file. The radius of the sphere is also changed as specified.

The completed model can then be rendered as an image, or exported from Maya as a 3D geometry file (e.g. STL, obj, etc).
